# Coexistence of phasmid sensory neurons and caudal glands offers a new perspective on cell type evolution in nematodes

**DOI:** 10.64898/2026.08.04.741185

**Authors:** Hyunsoo Yim, Ken C. Q. Nguyen, Luke T. Geiger, David H. Hall, Nathan E. Schroeder, Oliver Hobert

**Affiliations:** Columbia University, Department of Biological Sciences, Howard Hughes Medical Institute, New York, USA; Albert Einstein College of Medicine, Department of Neuroscience, New York, USA; Queens College CUNY, Department of Biology, New York, NY; University of Illinois at Urbana-Champaign, Department of Crop Science, Illinois, USA

## Abstract

The highly conserved body plan of nematodes makes members of this phylum excellent models to study cell type evolution. Early branching nematode lineages, mostly occupying aquatic habitats, usually contain caudal glands deployed for underwater attachment to a substrate, but have been thought to lack phasmid sensory organs, resulting in their historical classification as “Aphasmidia”. With the transition to a terrestrial environment, nematodes lost caudal glands and gained phasmid sensory neurons. The supposed mutually exclusive existence of caudal glands and phasmids has led to the suggestion that phasmid neurons may have evolved from caudal glands. Here, we rule out this possibility through light and electron microscopical analysis of *Mononchus aquaticus*, a member of the early branching Dorylaimia lineage, showing that phasmid sensory neurons and caudal glands do coexist. This observation not only argues against a proposed cell type evolution scenario accompanying aquatic-to-terrestrial transitions but also indicates that the presence of phasmid sensory organs may have been an ancestral trait of the nematode phylum.

## Text

In 1933, Chitwood proposed to group nematodes into two classes, the Phasmidia (also called Secernentea) and the earlier branching, more “basal” Aphasmidia (also called Adenophorea)^1^. This classification scheme persisted until around the turn of the century, when molecular phylogenetics showed that the Aphasmidia are not a monophyletic group ^2^. Nevertheless, the taxa previously subsumed under the Aphasmidia retained their position as early diverging lineages in nematode phylogeny.

There is a general consensus that nematodes are of a marine origin and that the earliest members contained a phylum-defining trait in the form of a set of caudal gland cells ^3,4^. Caudal glands are a characteristic feature of nematodes adapted to an aquatic environment, enabling animals to attach to a substratum underwater, using a presently ill-defined glue synthesized by the caudal glands and oftentimes released through a spinneret ^3,4^. Another feature of early branching nematode lineages is the presence of a nerve net of sensory neurons distributed throughout the body wall, yet - as the original term “Aphasmidia” indicated - these early branching lineages were never observed to contain phasmid sensory neurons in the tail of the animal ^3,4^. With the move to (semi-) terrestrial environments, nematodes were then thought to have lost caudal glands and gained phasmid neurons, leading to the suggestion that these neurons evolved from caudal glands ^3,4^.

*Mononchus aquaticus*, a voracious predator living in diverse wet habitats, is a member of the early branching Dorylaimia lineage, all members of which were previously considered to be part of the caudal gland-containing, phasmid-less Aphasmidia ^5^. Using scanning electron microscopy, we visualized features of *M. aquaticus* that are characteristic of early branching lineages (**Figure 1A-D**). At the anterior end of the animal, we noticed an elaborate amphid opening that is located posteriorly to the labial papilla, a characteristic feature of early branching nematodes (**Figure 1B**). We observed small cuticular openings distributed throughout the body wall (**Figure 1C**) whose number closely corresponds to the number of previously reported dye filling, body wall neurons of *M. aquaticus*, thus corroborating the presence of an extensive sensory nerve net with direct exposure to the environment ^6^. At the posterior end, a spinneret structure is apparent (**Figure 1D**) through which aquatic nematodes secrete a glue-like substance for anchoring to a substrate. We observe such anchoring behavior in *M. aquaticus* at all larval and adult stages. Caudal glands were also easily detectable by differential interference contrast microscopy (**Figure 1E**).

**Figure 1:**
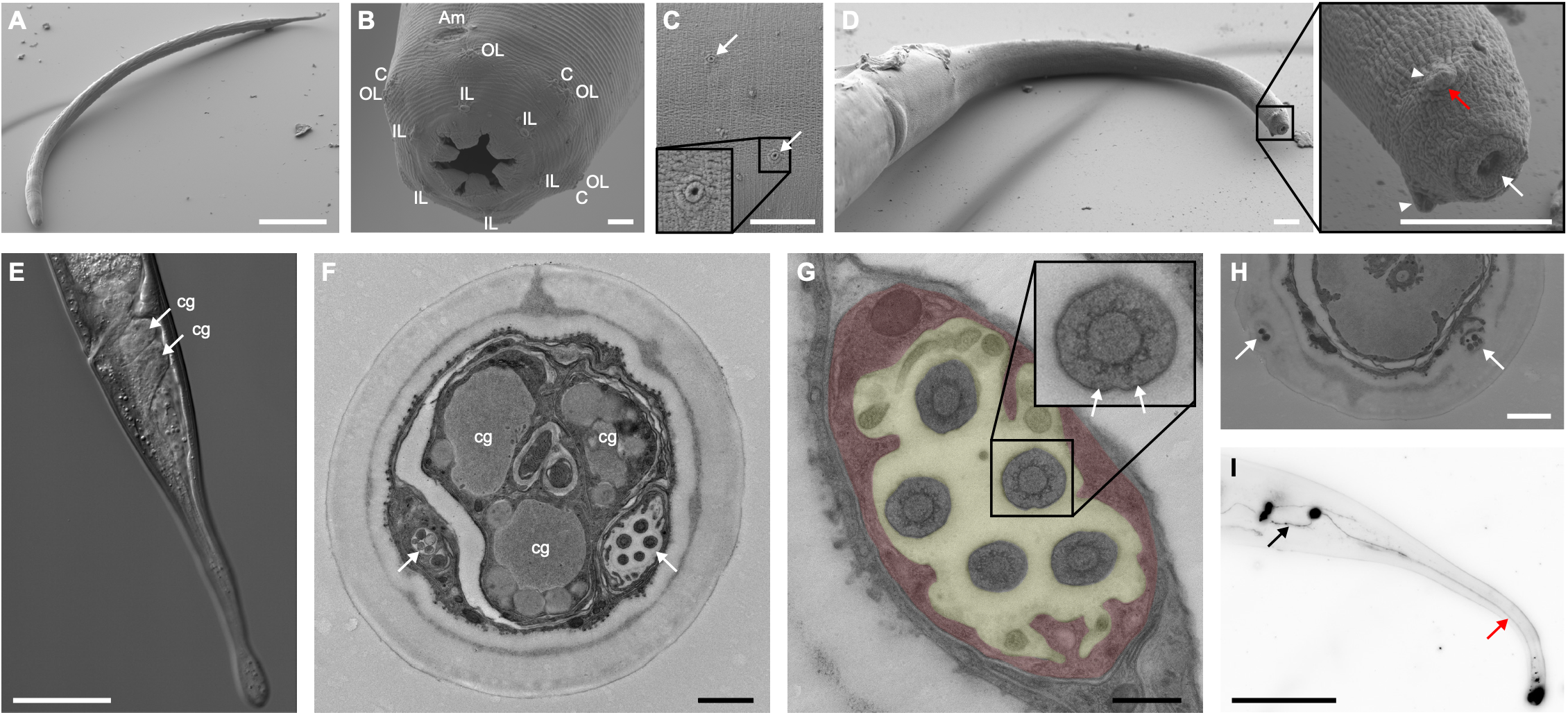
Anatomy of *Mononchus aquaticus*. **(A–D)** Scanning electron micrographs (SEM). **(A)** Low-magnification view of the whole animal. **(B)** Sensory structures of the head; Am, Amphid; C, Cephalic; OL: Outer labial; IL, Inner labial sensilla. **(C)** Small cuticular openings (arrow) distributed along the body wall. **(D)** High-magnification view of the tail terminus showing the coexistence of the spinneret (arrow) and the phasmidial papillae (arrowhead). The red arrow indicates the phasmid pore. **(E)** Nomarski DIC micrograph confirming the presence of the caudal glands. White arrows indicate the cell bodies of two caudal glands (cg). The third cell body is located in a different focal plane. (F–H) Transmission electron micrographs (TEM) of the tail. **(F)** Transverse section of the tail showing the phasmid sensillum (arrow) adjacent to the caudal gland ducts (cg). **(G)** Sensory cilia of the phasmid neurons showing ciliated endings with 9+0 microtubule doublet arrangements and Y-links (arrow). Red overlay highlights the phasmid sheath glia, and yellow overlay indicates the phasmid sheath lumen. **(H)** Phasmid opening at the phasmidial papilla. The arrow indicates ciliated endings of the phasmid neurons. **(I)** Fluorescence micrograph of a dye-filled (DiD) phasmid neuron. The black arrow indicates the anteriorly projecting axon, and the red arrow indicates the dendrite originating from the phasmidial papilla. Scale bars: A, 100 µm; B, 2 µm; C, 5 µm; D, 5 µm; E, 50 µm; F, 1 µm; G, 200 nm; H, 1 µm; I, 50 µm.

We were surprised to notice two cuticular protrusions close to the spinneret apparatus that are reminiscent of the phasmid sensory structures (**Figure 1D**). Since there has never been a report of a definitive coexistence of caudal glands and phasmids – which would have questioned the original “Aphasmidia” classification scheme – we took electron micrographs of transverse sections of the tail. This TEM analysis provided unequivocal evidence of the presence of phasmid neurons in *Mononchus aquaticus* since it revealed: (a) a clear phasmid opening with five ciliated endings exiting to the outside; (b) a characteristic 9+0 doublet microtubule configuration; (c) a classic Y-link images of a typical transition zone in a ciliated neuron; (d) surrounding sheath and socket glia cells that are characteristic of phasmid structures in Phasmidia ^7^ (**Figure 1F-H**). We captured the entire structure of the phasmid sensory neurons through dye filling, which revealed dendritic endings that extend to the same place where we observed the phasmid opening to be located. Moreover, we detected axons that extend anteriorly, possibly into the lumbar ganglion of the animals (**Figure 1I**).

These observations argue strongly against a classic model, most recently reformulated by Tahseen and Sudhaus in 2025 that was based on the presumed mutually exclusive existence of caudal glands and phasmid neurons ^4^. In this model, caudal glands are the precursor to the phasmid sensory neurons, thereby resolving the phasmid origin question, a question that Chitwood and Chitwood described as “one of the most perplexing problems in nematology” ^3^. This very attractive hypothesis, even though not universally accepted ^8^, is in line with more recent work in animal cell type evolution which has suggested secretory gland-like cells to be a potential precursor of neurons ^9^. The loss of a feature characteristic of aquatic nematodes (i.e. gland cells controlling an important function in an aqueous environment) and gain of a feature of terrestrial animals (i.e. chemosensory phasmid sensory neurons that control navigation) would have also presented a potential single cell paradigm for one of the most fascinating events in evolutionary biology, the aquatic to land transition. However, our finding of a coexistence of caudal glands and phasmid neurons disproves this scenario and rather indicates that the phasmids – together with the amphids – are a shared, phylum-defining trait.

Even though our high-resolution microscopical analysis ruled out a compelling scenario for the evolution of a specific cell type, we envision that a continued, detailed comparative anatomical analysis of diverse nematode species will suggest other fascinating evolutionary processes on the level of single cells.

## SUPPLEMENTAL INFORMATION

Supplemental information contains experimental procedures and can be found with this article online.

## DECLARATION OF INTERESTS

The authors declare no competing interests.

## Supplemental Information

### EXPERIMENTRAL PROCEDURES

#### Strain

The *M. aquaticus* culture used was initially isolated from a maize field at the Northwestern Illinois Agricultural Research and Demonstration Center in Monmouth, Illinois, as described before ^1^. Culture conditions were also as described before ^1^, with some modifications of the protocol that we will describe elsewhere.

#### Scanning Electron Microscopy (SEM)

For scanning electron microscopy, Monoicous aquaticus were fixed in protocol described in ^2^. Specimens were imaged using a Zeiss Supra 40 scanning electron microscope.

#### Transmission Electron Microscopy (TEM)

For TEM, we utilized a high-pressure freezing and freeze-substitution (HPF-FS) protocol described in ^3^, with a minor modification in the fixative composition. Specifically, we used a modified freeze-substitution fixative consisting of 3% osmium tetroxide, 0.2% uranyl acetate, and 2% double-distilled water in acetone.

#### Dye filling

We used a previous described protocol ^1^.

## Notes

### Competing Interest Statement

The authors have declared no competing interest.

### Funding

- Howard Hughes Medical Institute

## REFERENCES

1. Chitwood, B.G. (1933). Multiple Origins of Neurons From Secretory Cells. J. Parasitology 20, 131.

2. Blaxter, M.L., De Ley, P., Garey, J.R., Liu, L.X., Scheldeman, P., Vierstraete, A., Vanfleteren, J.R., Mackey, L.Y., Dorris, M., Frisse, L.M., et al. (1998). A molecular evolutionary framework for the phylum Nematoda. Nature 392, 71–75. 10.1038/32160.

3. Chitwood, B.G., and Chitwood, M.B. (1950). Introduction to Nematology (University Park Press).

4. Tahseen, Q., and Sudhaus, W. (2025). Caudal Glands in Nematodes: Morphology, Evolutionary Shifts and Functional Implications. Int. J. Zoo. Animal Biol. 8, 000654.

5. Ahmad, W., and Jairajpuri, M.S. Mononchida (2010) The Predatory Soil Nematodes (Brill).

6. Han, J., Ficca, A., Lanzatella, M., Leang, K., Barnum, M., Boudreaux, J.C.T., and Schroeder, N.E. (2026). Analysis of Nematode Ventral Nerve Cords Suggests Multiple Instances of Evolutionary Changes to Neuron Number. Evol Dev 28, e70037. 10.1111/ede.70037.

7. Hall, D.H., and Altun, Z. (2007). C. Elegans Atlas (Cold Spring Harbor Laboratory Press).

8. Lorenzen, S., and Lorenzen, S. (1994). The phylogenetic systematics of freeliving nematodes (Ray Society).

9. Moroz, L.L. (2021). Multiple Origins of Neurons From Secretory Cells. Front Cell Dev Biol 9, 669087. 10.3389/fcell.2021.669087.

## REFERENCES FOR SUPPLEMENTAL INFORMATION

1. Han, J., Ficca, A., Lanzatella, M., Leang, K., Barnum, M., Boudreaux, J.C.T., and Schroeder, N.E. (2026). Analysis of Nematode Ventral Nerve Cords Suggests Multiple Instances of Evolutionary Changes to Neuron Number. Evol Dev 28, e70037. 10.1111/ede.70037.

2. Hall, D.H., Winfrey, V.P., Blauer, G., Hoffman, L., Furuta, T., Rose, K.L., Hobert, O. and Greenstein, D. (1999). Ultrastructural Features of the Adult Hermaphrodite Gonad of Caenorhabditis elegans: Relations between the Germ Line and Soma. Dev. Biol. 212: 101–123. 10.1006/dbio.1999.9356

3. Hall, D.H, Hartwieg, E, and Nguyen, K.C.Q. (2012). Modern Electron Microscopy Methods for C. elegans. Methods in Cell Biol 107, 94–147. 10.1016/B978-0-12-394620-1.00004-7

